# Mitochondria define intestinal stem cell differentiation downstream of a FOXO/Notch axis

**DOI:** 10.1101/777391

**Authors:** M.C. Ludikhuize, M. Meerlo, M. Pages Gallego, M. Burgaya Julià, N.T.B. Nguyen, E. C. Brombacher, J.H. Paik, B.M. T. Burgering, M.J. Rodriguez Colman

## Abstract

Differential signalling of the WNT and Notch pathways regulates proliferation and differentiation of Lgr5^+^ crypt-based columnar cells (CBCs) into all cell lineages of the intestine. We have recently shown that high mitochondrial activity in CBCs is key in maintaining stem cell function. Interestingly, while high mitochondrial activity drives CBCs, it is reduced in the adjacent secretory Paneth cells (PCs). This observation implies that during differentiation towards PCs, CBCs undergo a metabolic rewiring involving downregulation of mitochondrial number and activity, through a hitherto unknown mechanism. Here we demonstrate, using intestinal organoids that FoxO transcription factors and Notch signalling functionally interact in determining CBC cell fate. In agreement with the organoid data, combined *Foxo1* and *3* deletion in mice increases PC number in the intestine. Importantly, we show that FOXO and Notch signalling converge onto regulation of mitochondrial fission, which in turn provokes stem cell differentiation into the secretory types; Goblet cells and PCs. Finally, mapping intestinal stem cell differentiation based on pseudotime computation of scRNA-seq data further supports the role of FOXO, Notch and mitochondria in determining secretory differentiation. This shows that mitochondria is not only a discriminatory hallmark of CBCs and PCs, but that its status actively determines lineage commitment during differentiation. Together, our work describes a new signalling-metabolic axis in stem cell differentiation and highlights the importance of mitochondria in determining cell fate.

## Introduction

The intestinal epithelium consists of multiple cell types that derive from one single precursor: the LGR5^+^ intestinal stem cell or crypt-based columnar cell (CBC) (Barker et al., 2007). WNT and Notch signalling pathways are the main regulators of CBCs maintenance, proliferation and differentiation. CBCs divide daily producing rapidly proliferating daughter cells known as transit amplifying cells (TA). Upon migration towards the intestinal lumen, progenitor cells become post-mitotic and can differentiate in two main lineages absorptive (enterocytes) or secretory types (goblet, Paneth, enteroendocrine and tuft cells) (reviewed in (Scoville et al., 2008)). Notch signalling is key in determining the choice between absorptive versus secretory lineages, in fact it acts as a node, while absorptive lineages sustain active Notch signalling, inactivation of Notch is a requisite for secretory cell differentiation (reviewed in Noah et al., 2011).

We have recently shown that CBCs and PCs, positioned adjacent in the intestinal crypt, are metabolically different (Rodríguez-Colman et al., 2017). While CBCs require mitochondrial respiration to maintain their stem cell function, Paneth cells are mainly glycolytic. Furthermore, mitochondrial metabolism appears to be reduced in the adjacent secretory Paneth cells. This observation implies that during differentiation towards Paneth cells, CBCs undergo a metabolic rewiring to downregulate of mitochondria. Importantly, there is a lack of understanding on how this process is regulated. Moreover, it remains unknown if there is a regulatory role of metabolic reprogramming per se in CBC differentiation.

Mitochondria have been shown to be involved in cell fate determination, i.e metabolic switches between glycolysis and OXPHOS accompanied by changes of mitochondrial shape are crucial for pluripotential reprogramming of induced pluripotent stem cells (Choi et al., 2015). Mitochondrial shape is regulated by ongoing cycles of fission and fusion (reviewed in Detmer and Chan, 2007). Importantly, these dynamics are important in the regulation of several cellular processes, i.e. cell cycle and apoptosis (Mitra, 2013; Pernas and Scorrano, 2016). Fission is executed by the GTPases DRP1 and DYN2, whereas the GTPases MFN1, MFN2, and OPA1 promote fusion. Upon activation DRP1 binds to adaptor proteins located in mitochondrial outer-membrane to initiate the process of fission. FIS1 is an adaptor protein that like DRP1 is conserved in yeast mice and human (Labbé et al., 2014) and both have been reported to be involved in stem cell maintenance and differentiation (reviewed in (Seo et al., 2018)).

The Forkhead box O (FOXO) transcription factors are regulated by insulin or growth factor-dependent PI3K signalling, glucose and oxidative stress. FOXOs have been reported to regulate maintenance of stemness in several systems, including embryonic (Zhang et al., 2011), hematopoietic (Miyamoto et al., 2007; Rimmelé et al., 2015) and neural stem cells (Paik et al., 2009; Renault et al., 2009). Mechanistically, FOXOs have been linked to numerous processes including cell cycle regulation, cell death, redox balance, mitochondrial metabolism, classical autophagy, but also mitophagy and mitochondrial fission through regulation of Fis1 (Rimmelé et al., 2015; Wang et al., 2012; Yalcin et al., 2008). At present it is unclear whether and how these various FOXO regulated processes contribute to aforementioned role in stemness regulation, although most studies invoke an ill-defined role for “oxidative stress”. FOXO interacts with several signalling pathways shown to be important in stem cell regulation including WNT (Eijkelenboom and Burgering, 2013) and Notch signalling pathways (Kitamura et al., 2007). In the mouse intestine it has been shown that FoxO1 loss induces differentiation of endocrine progenitors into glucose-responsive insulin-producing cells (Bouchi et al., 2014).

Here, we describe a novel function of FOXO as a regulator of intestinal homeostasis. We show that in organoids as well as in mice, conditional loss of FOXO leads to increase in the number of Paneth cells with a concomitant loss of LGR5+ cycling stem cells. This indicates a role of FOXO and PI3K signalling in the regulation of stem cell fate towards PCs. We describe that FOXO downregulation-driven secretory differentiation occurs along with Notch inhibition and conversely, Notch inhibition leads to decreased FOXO levels. Mechanistically we show that decreased mitochondrial respiration and increased mitochondrial fission, triggered by transcriptional regulation of miRNA 484 and upregulation of FIS1, are requisites for Paneth cell differentiation. Finally, *in silico* analysis of scRNA-seq supported our experimental data and further showed that both secretory types, Paneth and Goblet cells, are regulated by the Notch/FOXO/mitochondria axis. Together these results provide a comprehensive model in which classical cell signalling combines with mitochondrial metabolism, employing mitochondria as a signalling hub and as an essential regulator of stem cell maintenance and differentiation.

## Results

### Foxo1/3 down regulation induces secretory differentiation in vitro and in vivo

In order to analyse the role of FOXO in the intestine we introduced into normal small intestinal (SI) organoids doxycycline inducible short hairpin constructs to knockdown *Foxo1* and *3* expression (Charitou et al., 2015; Hornsveld et al., 2018). We confirmed FOXO1 and 3 knockdown by gene expression and protein level after 36 hrs of doxycycline treatment (Suppl. Fig. 1a and b). First, we analysed the effect of *Foxo* knockdown (KD) on the expression of intestinal cell type markers. *Foxo* KD resulted in specific downregulation of the stem cell markers *Lgr5*, *Ascl2* and *Olfm4* (Fig. 1a), as no doxycycline effect was observed in the Luciferase short hairpin control organoid line (Suppl. Fig. 1c). SI stem cells are proliferative and in agreement, expression of the proliferation marker *Ki67* was reduced and we observed reduced organoid size upon *Foxo* KD (Fig. 1a and Suppl. Fig.1d). Decreased proliferation is a prerequisite for cell differentiation. Consequently, we analysed the expression of several differentiation markers. While we did not find differences in the expression of enterocyte or enteroendocrine markers (Alpi, Sct and Neurog3), we found a significant increase in the expression of the secretory cell markers *Lyz1* and *2*, *Muc2* and *Gob5* upon *Foxo* KD (Fig. 1a). These results suggest that FOXOs positively regulate stem cell state and that its downregulation leads to an increased differentiation towards the secretory phenotype. RNA expression is not always indicative for protein expression. Therefore, we analysed the protein levels of the PC marker Lysozyme (LYS) by western blot, and we visualized the number and localization of PCs by confocal microscopy and WGA (Wheat Germ Agglutinin)-staining of organoids. FOXO loss induced an increase in Lysozyme protein level and in the number of Paneth cells allocated at the bottom of the intestinal crypt (Fig1. b and c). In order to quantify Paneth cell induction by Foxo KD, we measured CD24^+^ SSC^high^-cells by flow cytometry. In line with gene and protein expression analysis, we observed an increase in the population of Paneth cells upon *Foxo* KD (Fig. 1d and Suppl. Fig. 1e), further supporting the notion that *Foxo* KD drives PC differentiation. In order to corroborate our *in vitro* observations *in vivo*, we made use of previously generated Villin-Cre; FoxO 1,3,4 ko mice. In line with our previous observations, FOXO1 staining in intestinal sections of WT mice showed nuclear localization in stem cells and TA cells and negative staining in Paneth cells and Goblet cells (Fig. 1e). Villin-Cre: FoxO 1,3,4 ko mice showed indeed FOXO1, 3 and 4 knock-out specifically in the epithelial cells of the intestinal lining (Fig. 1e). Importantly, we observed that knock-out of *Foxo* induced a drastic increase in the population of Lys^+^ Paneth cells (Fig. 1f). Moreover, we found a decrease in BrdU^+^ cells at the bottom of the intestinal crypts, which could be a consequence of the expansion of PCs. Of note, Brdu^+^ cells are still present in the TA compartment after *Foxo* knock out (Suppl. Fig. 1f). Altogether, organoids and mouse analysis, support the notion that *Foxo* downregulation leads to stem cell differentiation into secretory cells.

**Figure 1.**
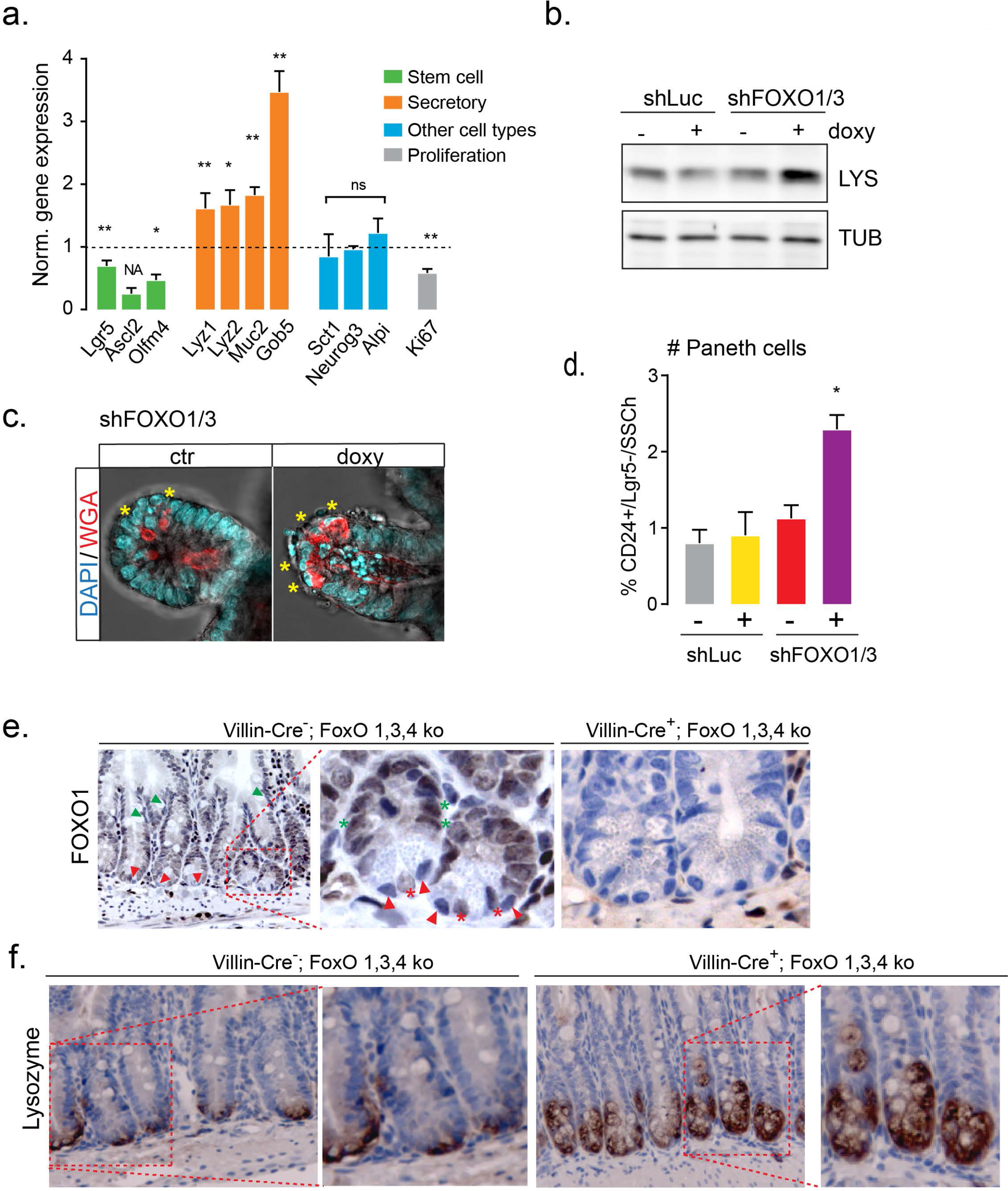
Foxo1/3 downregulation induces secretory differentiation in-vitro and in-vivo. **a.** qPCR gene expression analysis of intestinal cell lineages marker genes in doxycycline treated *shFoxo1/3* organoids. Bars represent the mean expression(± SEM, n=5) after normalization by non-treated organoids (see Fig. S1B for *shLuc* control organoids). **b.** Western blot detection of Lysozyme and Tubulin in *shLuc* and *shFoxo1/3* organoids (representative for n=4). **c.** Representative images of *shFoxo1 /3* organoids stained with DAPI and WGA for PC detection (asterisks indicate PCs). d.Paneth cell quantification by flow cytometry in DTR-Lgr5-GFP *shLuc* and *shFoxo1/3* organoids stained with anti-CD24. PCs: CD24^+^/LGR5^−^/SSC^high^ PCs (mean± SEM, n=4,) (for gating strategy see Fig. S1c). **e and f.** lmmunohistochemistry of crypts of Villlin-Cre-;Fox0 1,3,4L/L and Villin-Cre+ Fox0 1,3,4L/L mice. e. FOX01 staining in small intestine. Red arrowheads indicate Paneth cells, red asterisks indicate intestinal stem cells, green arrowheads indicate goblet cells and green asterisk indicate transit amplifying cells (TA). **f.** Lysozyme staining. In a. Dunn’s multiple comparisons test, ind Mann Whitney test test. *= p < 0.05, **= p<0.01, ***=p<0.001.

### Downregulation of Foxo1/3 leads to decreased mitochondrial respiration and induces mitochondrial fission through the regulation of miRNA-484

Our previous work demonstrated the importance of mitochondrial metabolism in CBCs. Moreover, we found that compared to stem cells, Paneth cells have lower mitochondrial activity (Rodríguez-Colman et al., 2017). This indicates that along differentiation from CBCs to Paneth cells, a metabolic reprogramming takes place in which mitochondrial number and activity are reduced. FoxO factors are well known regulators of metabolism (reviewed in (Eijkelenboom and Burgering, 2013)). Thus, to investigate the mechanisms that lead to Paneth cell differentiation upon FOXO loss, we analysed the metabolic consequences of *Foxo* KD in small intestinal organoids. Bioenergetics analysis showed that *Foxo* KD leads to decreased basal and maximal mitochondrial respiration (Fig. 2a and Suppl. Fig. 2a), and a milder decrease in basal glycolysis (Suppl. Fig. 2a and b). These observations indicate that FOXO loss affects mitochondrial function. Impairment of FOXO activity has been reported to affect the oxidative stress response (Essers et al., 2004; Ferber et al., 2012). In order to analyse the redox state in mitochondria, we introduced a genetically encoded probe (mtGrx1-roGFP2) in organoids to measure the ratio GSH/GSSG by confocal live imaging (Gutscher et al., 2008; Wong et al., 2017). Our results showed no significant changes in the mitochondrial redox state upon *Foxo* loss (Suppl. Fig. 2c). However, we found increased phosphorylation of the mitogen-activated protein kinase (MAPK) p38 upon *Foxo* KD (Suppl. Fig 2d), which is an indicator of cellular stress (Cuadrado and Nebreda, 2010).

**Figure 2.**
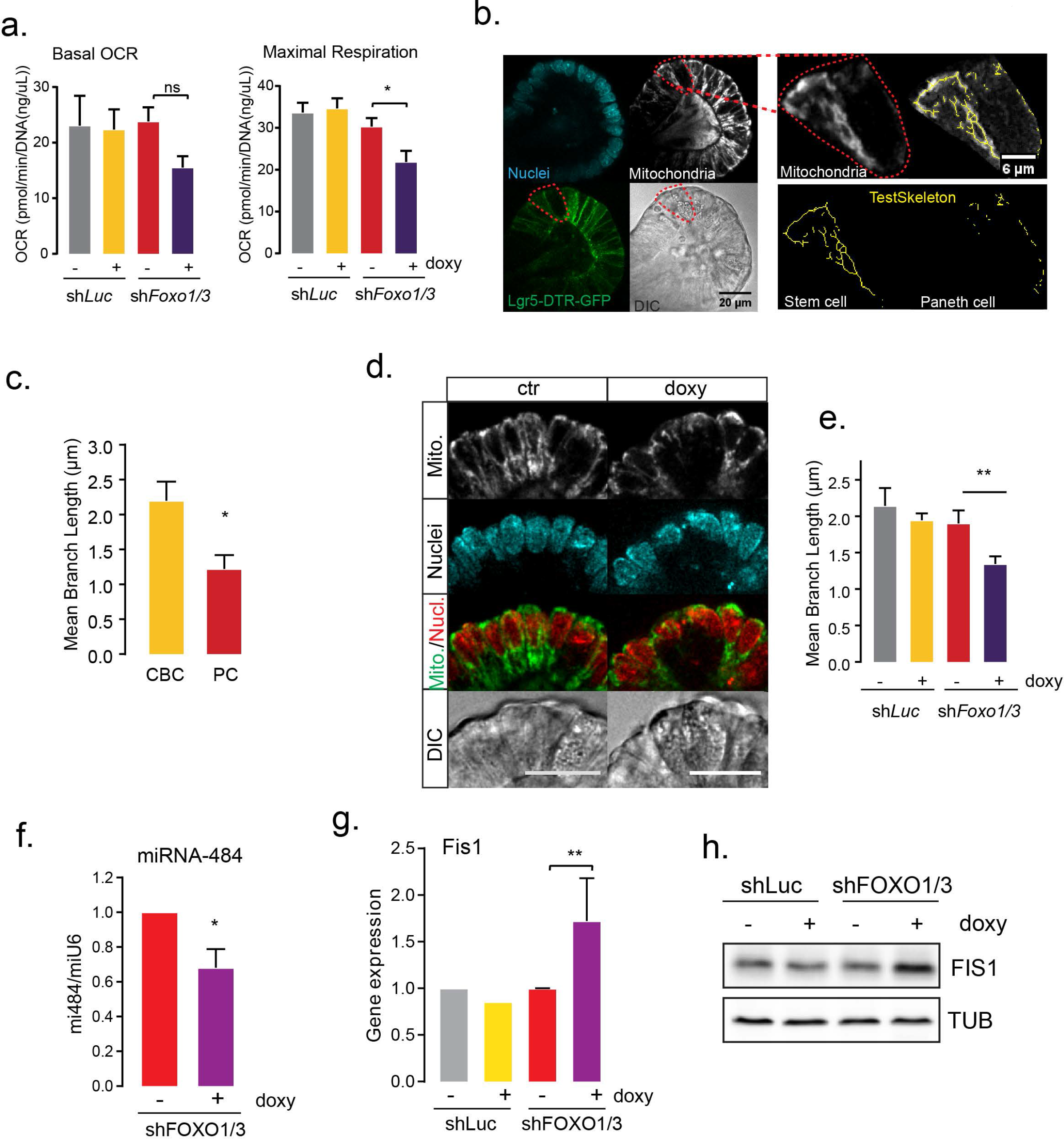
Downregulation of Foxo1/3 leads to decreased respiration and induces mitochondrial fission through the regulation of miRNA-484. **a.** Basal oxygen consumption rate (OCR) and maximal respiration of shLuc and *shFOX01/3* organoids were determined in mitochondrial stress test by Seahorse analysis (mean ± SEM, n=3). **b.** Representative images of DTR-Lgr5-GFP organoids stained with Hoechst and Mitotracker **c.** Quantification of mitochondrial mean branch length in CBCs and PCs (mean ± SEM n= 6 PC and 8 CBCs in 2 independent experiments, T-test). **d.** Representative images of doxycycline-treated *shFoxo1/3* organoids stained with Mitotracker. **e.** Quantification of mitochondrial mean branch length of shLuc and *shFoxo1/3* organoids (mean± SEM, n=6 and 7 cells per condition). **f.** Relative expression of miRNA-484 in doxycycline treated *shFoxo1/3* organoids determined by qPCR (mean± SEM, n=5, Student’s T-test). **g.** Relative gene expression of Fis1 in shLuc and *shFoxo1/3* organoids determined by qPCR (mean± SD, n=6, Mann-Whitney test). **h.** Western blot analysis of FIS1in shLuc and *shFoxo1/3* organoids 10 (representative for n=4). In a. and e. Holm-Sidak’s multiple comparisons test.*= p < 0.05, **= p<0.01.

To further study the consequence of FOXO loss in mitochondria, we analysed the morphology of mitochondria in small intestinal organoids. Mitochondria are dynamic organelles that are maintained by cycles of fission and fusion. Fission and fusion rates determine the morphology of mitochondria, from more fragmented to more elongated structures respectively. First, we analysed mitochondria and their morphology in CBCs and PCs. We observed decreased amount of mitochondria in PCs when compared to adjacent stem cells (Fig. 2b). In addition to that we found morphological differences, while stem cells showed fragmented and clearly distinguishable fused mitochondria, Paneth cells lacked of fused mitochondrial structures and appeared mainly fragmented (Fig. 2b and c). Next, we analysed mitochondrial morphology upon *Foxo* knockdown. We found that while in control organoids mitochondria display both morphologies, fused and fragmented, *Foxo* loss leads to decreased fused structures and an enrichment in fragmented mitochondria (Fig. 2d and e). Together, our results show that FOXO loss leads to decreased mitochondrial respiration, increased mitochondrial fragmentation. Next, we investigated the mechanisms by which FOXO drives changes in mitochondria. The balance between mitochondrial fission and fusion is maintained by the activity of a set of dynamin-related GTPases that drive these processes. Recruitment of the GTPase DRP1 to mitochondria is a critical step in fission. Fis1p recruits the Drp1P homolog to mitochondria in yeast, and in mouse and human its knockdown induces elongation, and its overexpression increases fragmentation in a cell type specific manner (Losón et al., 2013; Pernas and Scorrano, 2016; Yu et al., 2019). FOXO3 can regulate FIS1 levels through miR-484-mediated transcriptional regulation of Fis1 (Wang et al., 2012). Therefore, we analysed the effect of *Foxo* KD on miR-484, *Fis1* mRNA and FIS1 protein. We found that downregulation of FOXO resulted in a decrease in miR-484 expression and an increase *Fis1* mRNA and protein levels (Fig. 2 f-h). This suggests that FOXO regulation of *Fis1* may proceed through miR-484 regulation and suggests mitochondrial fission as a driver of PC differentiation upon FOXO downregulation.

### Mitochondrial fission is a requisite for PC differentiation

In order to assess if mitochondrial fission drives differentiation into Paneth cells upon *Foxo* KD, we made use of Mdivi-1 a small molecule inhibitor of mitochondrial fission. The suggested mode of action of Mdivi-1 is to inhibit DRP1 and it has been extensively used for inhibition of mitochondrial fission (Reddy, 2014; Smith and Gallo, 2017). First, we treated *shFoxo1/3* organoids with Mdivi-1 and analysed its effect in mitochondrial morphology by confocal microscopy. We found that Mdivi-1 was able to partially restore the increased fragmentation of mitochondria induced by FOXO loss (Fig. 3a and b). Then, we analysed the effect of inhibition of mitochondrial fission on Paneth cell differentiation. We analysed the gene expression of differentiation markers and protein levels of the PC marker Lysozyme. Gene expression analysis showed that although Mdivi-1 has no significant effect on the expression of stem cell markers upon FOXO loss, it does partially revert the gene expression induction of secretory markers (Suppl. Fig. 3a). Supporting this observation, we found that FOXO loss induction of the Paneth cell marker Lysozyme is rescued by inhibition of mitochondrial fission by Mdivi-1 (Suppl. Fig. 3b). Moreover, we found that inhibition of mitochondrial fission by Mdivi-1, can partially revert the increase in number of Paneth cells induced by FOXO loss (Fig. 3c and Suppl. Fig. 3f). Mdvi-1 treatment was not able to significantly recover mitochondrial respiration upon *Foxo* KD (Suppl. Fig. 3c). This suggests that possibly both, morphology and activity of mitochondria, are required for full reversion of the FOXO-loss induced phenotype. Altogether our results indicate that mitochondrial fission is a required step in the process of secretory differentiation induced by FOXO loss.

**Figure 3.**
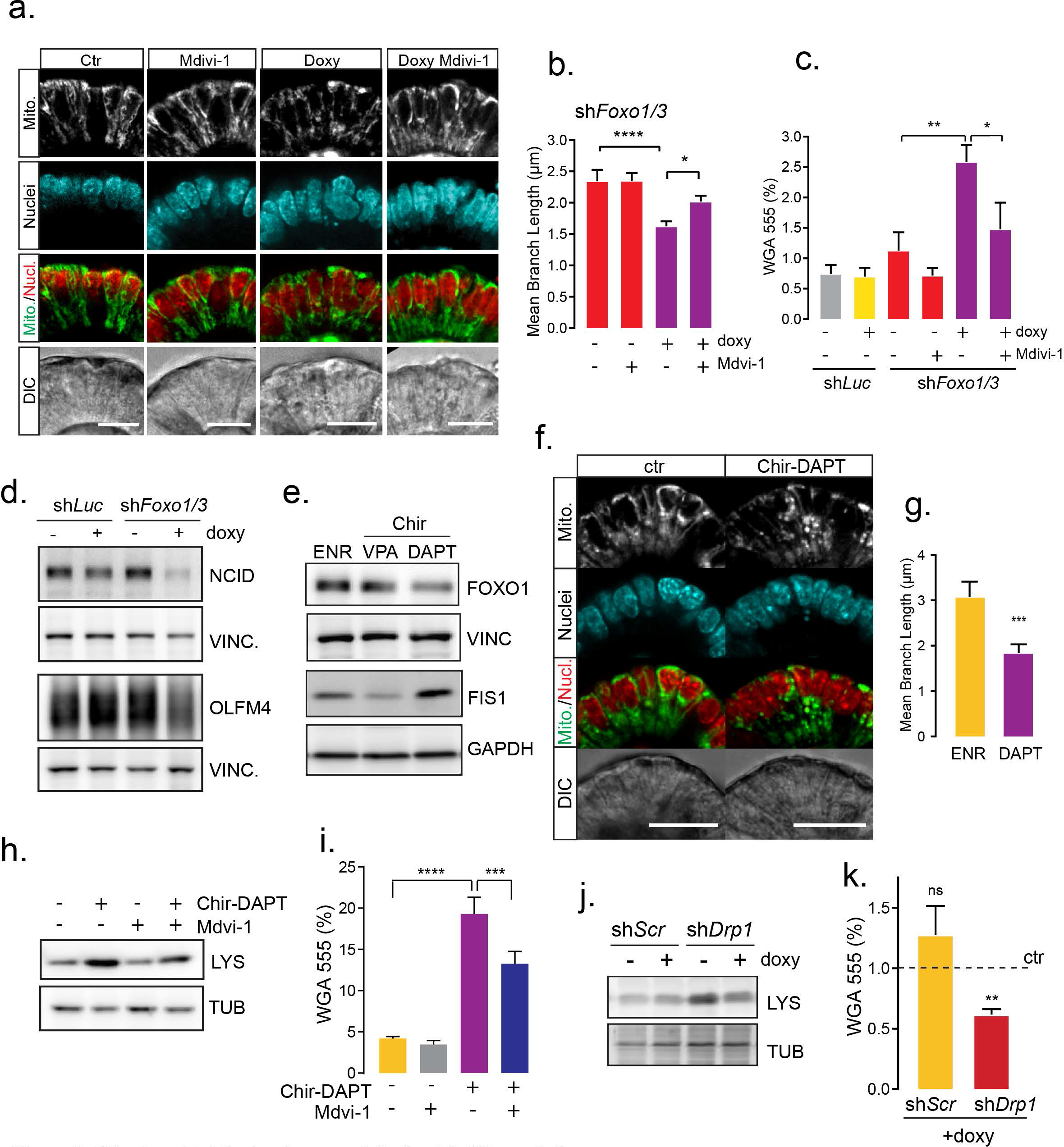
Mitochondrial fission is a requisite for PC differentiation. **a.** Representative images of *shFoxo1/3* organoids treated with doxycycline and Mdivi-1 and stained with Hoechst and Mitotracker (scale bar= 25 um, representative for n=3). **b.** Quantification of mitochondrial mean branch length (mean ± SEM, minimum of 20 cells from 3 independent experiments, Dunn’s multiple comparisons test). c. PC quantification by flow cytometry of DTR-LgrS-GFP shLuc and *shFoxo1/3* organoids treated with doxycycline and Mdivi-1 and stained with WGA-TMRE to determine percentage of WGN/LgrS-PCs (mean± SEM, test, n=S, Sidak’s multiple comparisons test). **d.** NCID and OLFM4 levels in shLuc and *shFoxo1/3* organoids by western blot (representative of n=4). **f.** FOX01 and FIS1 analyzed by western blot. ENR control, DAPT: PC enriched, VPA: Stem cell enriched organoids. Organoids were grown for 48 hs in normal medium (ENR) or with the addition of CHIR9921 and DAPT or CHIR and VPA (representative for n=3). **f.** Representative images of control and PC enriched organoids with Hoechst and Mitotracker. **g.** Quantification of mitochondrial mean branch length (mean ±SEM, minimum of 12 cells per condition from 3 independent experiments, Mann Whitney test) Scale bar 20um. **h and j.** Western blot analysis of Lysozyme and tubulin (representative of n=3 and n=2 respectively). **i and k.** PC analysis by WGA staining and flow cytometry of DTR-LgrS-GFP organoids treated with CHIR-DAPT and Mdivi-1 (n=7, mean ±SEM, Sidak’s multiple comparisons test) and DTR-LgrS-GFP shScr and *sh0rp1* organoids treated with doxycycline for 48 hs. Data is represented as ratio to non-treated shScr and *sh0rp1* organoids 12 (mean ± SD, n=4, Student’s T-test). *= p < 0.05, **= p<0.01, ***=p<0.001.

The intestinal stem cell state is maintained by high activity of WNT and Notch signalling pathways, and inhibition of Notch drives Paneth cell differentiation. FOXO and Notch have been reported to functionally interact in the regulation of stem cell maintenance and differentiation in muscle (Kitamura et al., 2007). Thus, we analysed the activation of Notch signalling upon Foxo knockdown. We observed that activated Notch (Notch Intracellular Cytoplasmatic Domain (NCID)) levels decreased upon Foxo KD (Fig. 3d). In agreement, the downstream target of Notch, OLFM4, is downregulated upon Foxo knockdown (Fig. 3d). This showed that PC differentiation induced by Foxo KD correlates with Notch inhibition. Organoid cultures can be customized to induce differentiation to specific cell types (Yin et al., 2014). In order to analyse the role of Foxo in canonical PC differentiation, we grew PC enriched organoid cultures by addition of DAPT and Chir9902 (Notch inhibition and WNT activation respectively) (Suppl. Fig. 3d). Additionally, we induced the formation of stem cell enriched cultures by a combination of Chir99021 and Valproic acid (VPA). Interestingly, FOXO levels seem to be responsive to Notch activity, as the induction of Paneth cell formation by Notch inhibition leads to a concomitant decrease in FOXO1 protein (Fig. 3e). In order to explore this further, we analysed the consequences of Notch inhibition on mitochondria. Similarly, to Foxo KD, we found that DAPT-mediated Notch inhibition induced the levels of FIS1 and mitochondrial fragmentation (Fig. 3e-g). Additionally, we observed that while mitochondrial DNA copy number is induced in stem cell enriched cultures, it is decreased in Paneth cell enriched cultures (Suppl. Fig. 3e). These results suggest that FOXO and Notch are mutually regulated and shows that inactivation of both pathways leads to FIS1 upregulation and mitochondrial fragmentation. Next, we analysed the effect of inhibition of mitochondrial fission on Notch inhibition-induced phenotypes by combining DAPT and Mdivi-1 treatments. Our results show that inhibition of mitochondrial fission by Mdivi-1 can partially revert LYS induction and the increase in PCs induced by Notch inhibition (Fig. 3h and i). These results indicate mitochondrial fission as a potential driver of differentiation. In order to analyse if manipulation of mitochondrial dynamics is sufficient to drive Paneth cell differentiation, we manipulated fission by introducing inducible short hairpin targeting the mitochondrial fission protein DRP1 (Suppl. Fig. 3g). Quantification of Paneth cells upon the knockdown of *Drp1* in basal conditions shows that *Drp1* loss resulted in a decrease in Lys protein and in the population of Paneth cells (Fig. 3j and k), supporting the notion that mitochondrial fission is a driver event of Paneth cell differentiation. Together our results show that in the intestine FOXO and Notch signalling converge to regulate stem cell maintenance, as inhibition of either FOXO Notch leads to Paneth cells differentiation through a mechanism that requires mitochondrial fission and downregulation.

### Small intestine sc-RNAseq analysis reveals a nodal role for Foxo/mitochondria in stem cell differentiation

In order to extend and independently confirm our understanding of the FOXO/Notch/mitochondria axis in the regulation of intestinal cell lineages, we re-analysed single cell sequencing data from mouse intestine (Haber et al., 2017). In the aforecited work, Haber et al performed full-length scRNA-seq of mouse small intestine of 8 mice with a total of 1522 cells. Clustering scRNA-seq data by t-Distributed Stochastic Neighbor Embedding (t-SNE) showed 9 cell clusters corresponding to the main cell lineages in mouse small intestine, in agreement with Haber et al. (Haber et al., 2017) (Fig. 4a and Suppl. Fig. 4a). In order to generate a temporal map of intestinal differentiation, we first applied dimension reduction and generated diffusion maps. We found that the top 2 diffusion components accommodate the cells in 3 branches. When plotting cluster classification to the diffusion components we found the stem cells located in the centre branching into 3 trajectories encompassing i) enterocytes ii) Tuft cells and iii) the secretory cluster of Goblet and Paneth cells on the same trajectory (Fig. 4b). This data representation indicates that enterocytes, Tuft and Goblet together with Paneth cells represent 3 main differentiation trajectories. Next we applied *Palantir* algorithm to estimate a pseudotime trajectory and branch probability through the data (Setty et al., 2018, 2019). Branch probability assigns to each cell a probability to differentiate into each potential terminal state. Thus, we transformed for the cluster classification of each cell from a discrete variable (i.e. observed in t-SNE plot) to a continuous one that more faithfully represents the process of differentiation (Suppl. Fig. 4b). Pseudotime calculation assigns to each cell a relative distance value to the starting stem cell state. Accordingly, the lowest values of pseudo-time were found in the center-located stem cells and these values increase along the branches in the direction of differentiation (Fig. 4b). We validated the performance of our pseudotime estimation by analysing branch probabilities and gene trajectories of the marker genes of each differentiation cluster. As expected, the expression of stem cell genes decreased along differentiation into all 3 differentiation trajectories, while the according differentiation markers specifically increased within each respective cluster (Fig. 4c). Of note, the expression of genes that define Goblet cell state also increased along pseudotime in the Paneth cell cluster and vice-versa, suggesting similarity in the trajectory of differentiation of these two cell types. Next, we analysed the expression of *Foxo1* and *Foxo3* in the diffusion components and along the differentiation trajectories. We found that along the progression to enterocytes or enteroendocrine differentiation the expression of *Foxo1* and *3* does not display major changes, however it decreases towards differentiation into Goblet, Paneth and to a lesser extent Tuft cells (Fig. 4d). Thus consistent with our experimental results, *Foxo1* and *Foxo3* are highly expressed in stem cells and downregulated in the secretory cell types (Suppl. Fig. 4c). As our experimental results showed that the regulation of stem cell maintenance and Paneth cell differentiation through FOXO occurs in a mitochondrial-dependent mechanism, we analysed the expression of genes that encode for mitochondrial proteins as a proxy of mitochondrial abundance. In agreement with our results, we found that mitochondria are high at low pseudotime (closer to the stem cell state) and decrease along differentiation trajectory into Paneth, Goblet cells and to lesser extent EE and Tuft cells, while it is maintained along enterocyte differentiation (Fig. 4d and Suppl. Fig. 4c). Lastly, we evaluated *Foxo1*, *Foxo3* and *Notch1* expression patterns in a cluster-independent manner. To do so, we defined the expression of the 3 genes as unique components to distribute all intestinal cells in a 3-dimensional space. Additionally, we introduced a fourth dimension by colouring each cell according their cluster classification, and alternatively by the expression level of the mitochondrial gene *Vdac1*. Importantly, this analysis shows that expression of these 3 genes, *Foxo1*, *Foxo3* and *Notch1* is sufficient to segregate the cells according to their lineages and secondly that decreased expression of *Foxo1*, *Foxo3*, *Notch1* and mitochondria are consistent and robust markers of Goblet and Paneth cell differentiation (Fig. 4e and Suppl. Fig. 4f). Lastly, we performed the same analysis but yet on another independent data set generated by droplet sc-RNAseq of 10396 cells of the small intestine (Haber et al., 2017) which lead to the same conclusions (Suppl. Fig. 4d-f). Thus taken together, this independent approach using sc-RNAseq analysis clearly confirmed our *in-vivo* and *in-vitro* observations; showing that FOXO and Notch signalling converging on mitochondria are key factors in intestinal stem cell maintenance and secretory cell differentiation.

**Figure 4.**
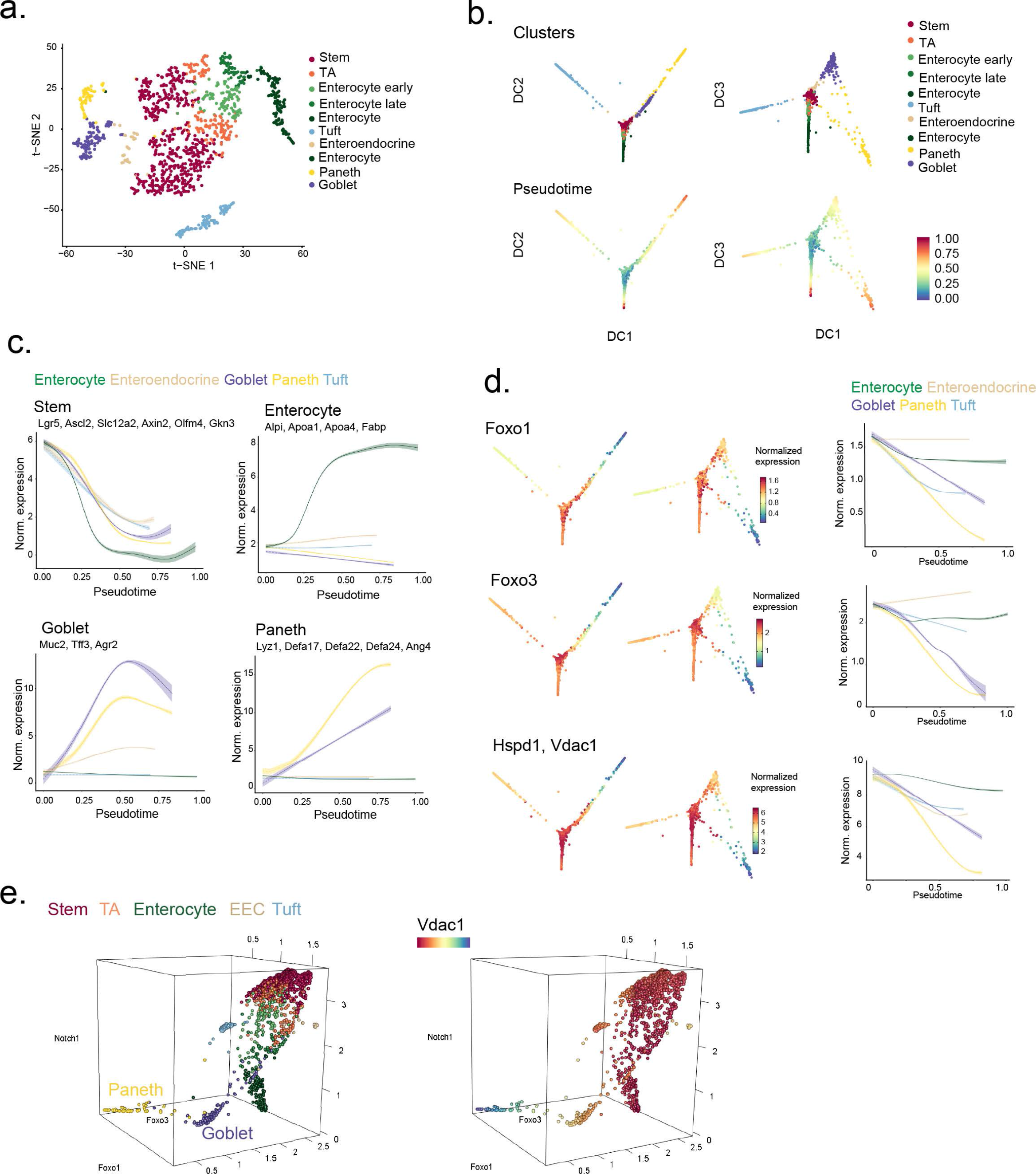
Small intestine sc-RNAseq analysis reveals a nodal role for Foxo and mitochondria in stem cell differentiation. **a.** t-SNE visualization of 1,522 single cells full-length sc-RNAseq data. Cluster annotation was done on the expression of known cell type markers (See Suppl. Fig. 6a). **b.** Representation of cell clusters and pseudo-time overlaid on the top 3 Diffusion Components (DC). DC were computed with destiny. Pseudo-time was calculated with Palantir. **c.** Validation of pseudo-time computation. Gene trends of cell type markers are plotted on pseudo-time for each cell cluster. **d.** Expression levels of Foxo1, 3 and mitochondrial markers (Hspd1 and Vdac) overlaid on the top 3 DC. Foxo1, 3 and mitochondrial gene trends are plotted on pseudo-time for each cell type cluster as a proxy of gene expression dynamics during cell differentiation. **e.** 3 dimensional representation of gene expression of Foxo1, Foxo3 and Notch1 in intestinal cells. As a fourth dimension cluster classification is assigned by color (right) or colored according Vdac1 expression levels (left).

## Discussion

WNT and Notch are evolutionary conserved signalling pathways that regulate cell patterning during development. Importantly in the mammalian intestine these same pathways control stem cell proliferation and differentiation. While WNT and Notch signalling are active in stem cells, in progenitor cells their activity is polarised, absorptive progenitors are WNT low and Notch high, whereas secretory progenitors are WNT high and Notch low (Sancho et al., 2015). In a recent study, we showed that mitochondrial metabolism is a key factor in CBCs maintenance and that CBCs and PCs metabolically interact (Rodríguez-Colman et al., 2017). In the present study we provide evidence that mitochondria play a key role in determining cell fate. Here we show that mitochondrial downregulation leads to stem cell differentiation into secretory lineages, while differentiation into the absorptive enterocyte lineage remains unchanged. Thus, mitochondrial function provides specificity in differentiation. The switch between mitochondria on and mitochondria off that we found driving differentiation coincides with the branching in Notch signalling between secretory and absorptive progenitors. Mechanistically, we show that, mitochondria and Notch, are interconnected through FoxO signalling. In contrast to WNT and Notch signalling, knowledge on the involvement of PI3K signalling in determining stem cell fate has been limited. Here we show that FoxO transcription factors, important downstream components in PI3K signalling, act as regulators of intestinal homeostasis. In fact, *Foxo1/3* silencing is sufficient to drive secretory fate commitment. Importantly, we found that the mechanism downstream FOXO involves regulation of FIS1 protein, through transcriptional regulation of miRNA-484, and consequently regulation of mitochondrial fission. Moreover, we found that *Foxo* downregulation leads to Notch inactivation, and conversely, Notch inhibition leads to decreased FOXO levels. Notch and FoxO signalling pathways have been shown to interact in other systems, in fact *Foxo1* ablation simulates *Notch1* ablation in mice (Hosaka et al., 2004; Krebs et al., 2000). Moreover, in muscle the Notch inhibitory effect on myoblast differentiation is FOXO1 dependent and conversely FOXO1 myogenesis is mediated through Notch interaction (Kitamura et al., 2007). In the same study it is shown that FOXO1 physically interacts with Notch and binds to the Csl DNA-binding protein to induce expression of the Notch target *Hes1*. In line with that, FOXO1 and Notch pathway functional interaction contributes to the inhibition of NSPC differentiation (Kim et al., 2015a). In a different study in skeletal muscle stem cells, it is shown that conditional deletion of *Foxo3* impairs stem self-renewal and increases differentiation through a mechanism that involves FOXO3 transcriptional regulation of *Notch1* and *Notch3* (Gopinath et al., 2014). Interestingly, FOXO transcriptional regulation of *Notch1* and *3* does not appear to be relevant in the small intestine as KD of *Foxo1/3* does not result in significant changes in *Notch1* expression (Suppl. Fig.4a). In our work, we show that FOXO regulates stem cell differentiation through a mitochondrial-dependent mechanism. However, FoxO factors have been involved in multiple cellular processes that potentially interact with mitochondrial function most notably oxidative stress and redox homeostasis (Essers et al., 2004; van Doeselaar and Burgering, 2018). In fact, the oxidative stress responsive MAPK p38 has been reported by us and others to be required for intestinal homeostasis (Houde et al., 2001; Rodríguez-Colman et al., 2017; Wakeman et al., 2012). Thus, in addition to mitochondrial fission, p38 could be involved in secretory differentiation upon *Foxo* knockdown. Indeed, we found p38 activation upon *Foxo* downregulation, however this activation did not appear to be triggered by mitochondrial ROS signalling as mitochondrial redox state was unperturbed upon FOXO loss. Mitochondrial fission has been previously linked to cell differentiation. Recently, FIS1-induced mitophagy has been reported to regulate stemness and differentiation in acute myeloid leukemia (Pei et al., 2018). In addition, DRP1-dependent mitochondrial fission is crucial for embryonic and cellular differentiation *in vivo* and *in vitro* (reviewed in Seo et al., 2018). In adult neural stem cells, DRP1 inhibition causes abnormal migration and prevents of neuronal differentiation and, in agreement with our observations, Midvi-1 treatment prevents neuronal differentiation (Kim et al., 2015b). Furthermore, during early myogenic differentiation DRP1-induced mitophagy is required (Chen et al., 2013) and Mdivi-1 treatment supresses the expression of myogenic regulatory factors, such as *MyoD* and *Myogenin* (Kim et al., 2013). Here, we showed that inhibition of mitochondrial fission by Mdivi-1 significantly inhibits the secretory phenotype induced upon *Foxo* KD or Notch inhibition, however it fails to recover the phenotype to the basal non-treated conditions. In this respect, it should be considered that Mdivi-1 can rescue the morphological phenotype of mitochondria but it fails to restore mitochondrial respiration. In addition, p38 is activated upon *Foxo* KD and can potentially contribute to secretory differentiation and Mdivi-1 treatment does not prevent p38 phosphorylation/activation upon *Foxo* KD (Suppl. Fig. 2d), suggesting a cooperative role for p38 and mitochondrial status. It would be interesting to further elucidate the role of mitochondrial respiration and p38 in differentiation. In this work we combine *in-vitro* and *in-vivo* data with *in-silico* analysis to deepen our understanding of the regulation of intestinal homeostasis. Previously published sc-RNAseq data on intestinal cells further and independently support our experimental data by showing that similar dynamics are observed for Notch/FOXO and mitochondria in the stem cell state and across differentiation into the differentiated lineages.

Additionally, we could discriminate between FoxO1 and 3 and found that both transcription factors behave comparably when analysing the stem cell state and differentiation into Goblet and Paneth cells. Furthermore, we found that while Foxo1 and 3 downregulation can define differentiation into Goblet and Paneth cells, other secretory types like Tuft cells differentiate independently of *Foxo1/3* downregulation. Importantly, Tuft cells branch separately from other lineages, suggesting that they are lowly related to these other cell types. These type of analysis provide additional and valuable information to better understand differentiation of relative less well studied lineages such as Tuft cells, for which the cell differentiation process is still a matter of debate (reviewed in Noah and Shroyer, 2013).

The balance between self-renewal and differentiation is under stringent control to maintain proper homeostasis. Uncontrolled growth can lead to intestinal hyperplasia, inflammatory processes and cancer. Our work highlights the importance of the cross-talk between cell signalling and metabolism, we show that this interaction regulates tissue homeostasis by defining cell fate and tissue heterogeneity. Notch signalling is one of the main signalling pathways that regulates tissue patterning and development and it has been recently shown that a glycolytic gradient also occurs during in organogenesis-stage mouse embryos (Bulusu et al., 2017). It would be of special interest to relate Notch signalling dynamics during development with changes in mitochondrial activation. These dynamics in development between signalling and metabolism are likely to be re-iterated upon tissue regeneration and possibly even in tumorigenesis. Non-genetic or phenotypic tumour heterogeneity is an emerging field of research, which can explain the chemotherapy surveillance, tumour regrowth and relapse in cases of genetic homogeneity (Roerink et al., 2018). Aberrant cell signalling connecting to metabolism rewiring may drive non-genetic heterogeneity and better understanding of these processes can contribute to improved cancer therapy as well as understanding fundamental biology.

---

**Suppl. Figure 1:**
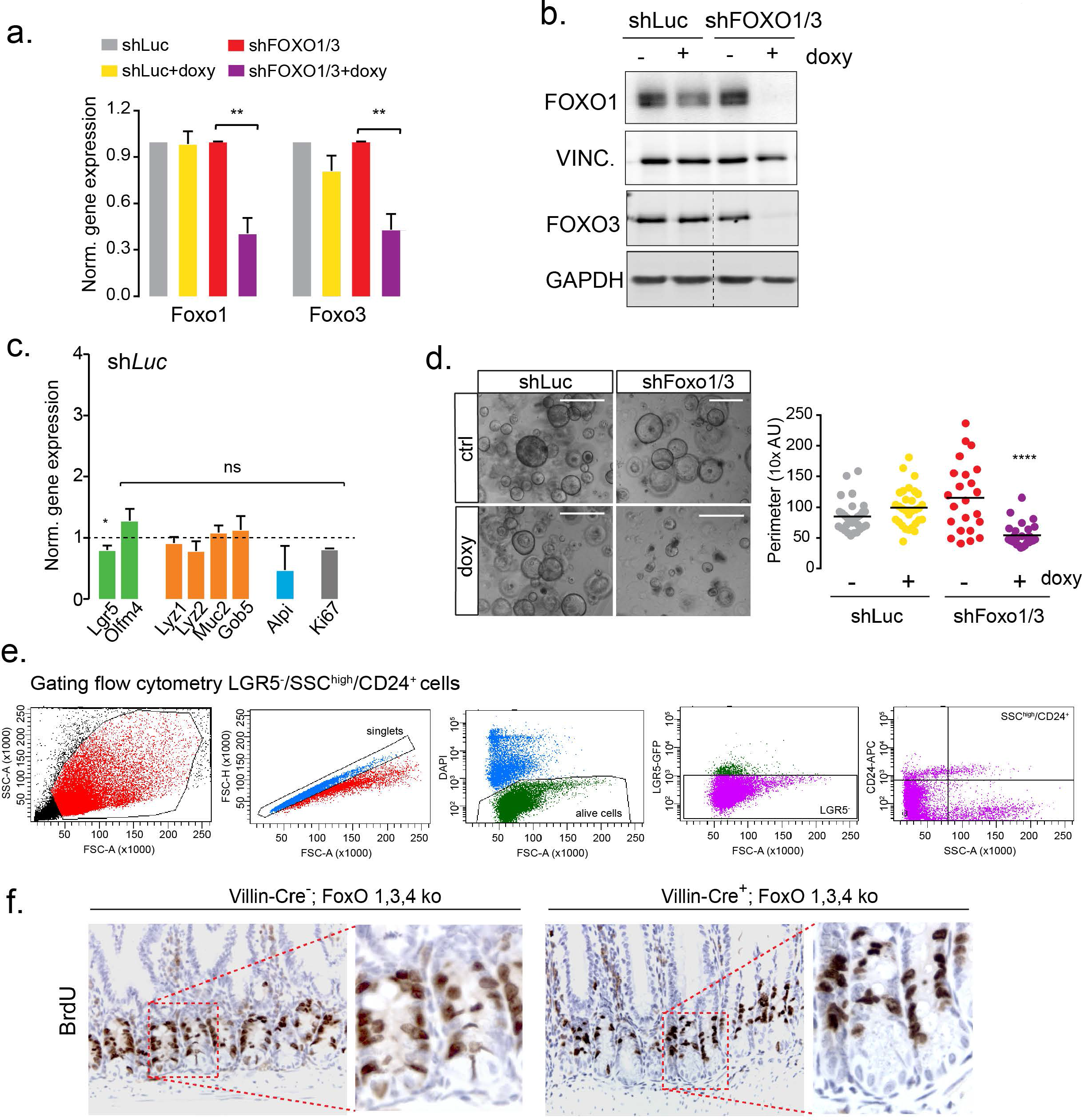
Foxo1/3 downregulation induces secretory differentiation in-vitro and in-vivo. **a.** Foxo1 and Foxo3 expression in doxycycline treated shLuc and shFoxo1/3 organoids by qPCR (mean± SEM, n=4) (see also Fig. S1a for protein expression). **b.** Western blot analysis of shluc and shFoxo1/3 organoids, VINCULIN and GAPDH are used as loading controls (representative for n=3). **c.** Gene expression analysis of shLuc organoids by qPCR and normalized by the non-treated condition (mean± SEM, n=3-5. ANOVA and Dunns multiple comparison test). **d.** Representative images of doxycycline treated shLuc and shFoxo1/3 organoids cultured in WENR medium (scale bar= 500 AU) and quantification of their perimeter (one representative experiment of 3, each dot represents one organoid, ANOVA and Dunn’s multiple comparison test) **e.** Flow cytometry gating strategy for quantification of live PCs. **f.** lmmunohistochemistry of crypts of Villlin-g Cre-;Fox01,3,4L/L and Villin-Cre+ Fox0 1,3,4L/L mice. Proliferative cells are marked by BrdU positive staining.

**Figure S2.**
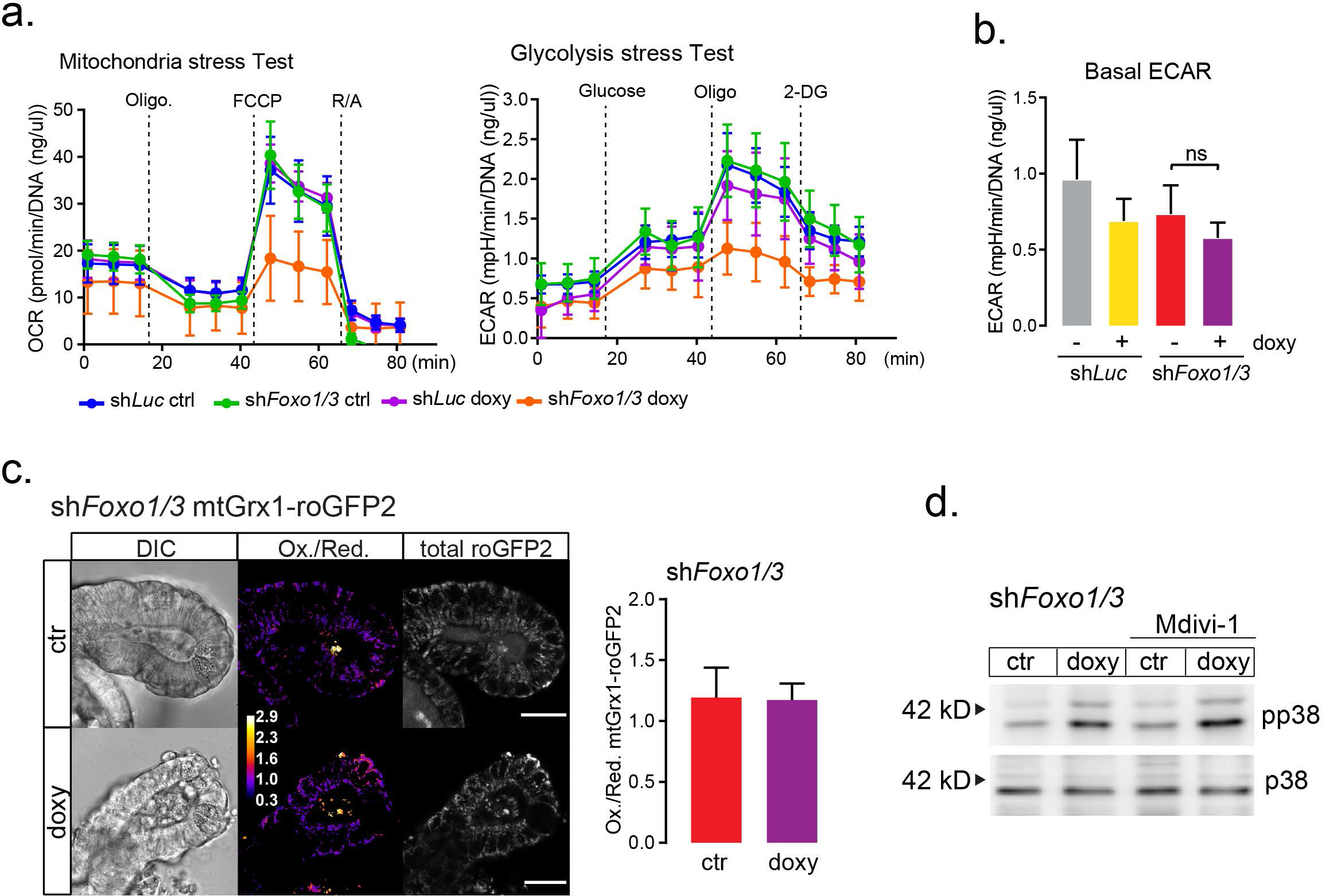
Foxo1/3 downregulation has a mild effect Glycolysis, no affect in the mitochondrial redox state, but induces p38 phosphorylation. **A and b.** Representative Mitochondria stress test and Glycolytic stress test of shLuc and *shFoxo1/3* organoids measured with Seahorse Analyzer. **c.** Basal ECAR of shLuc and *shFoxo1/3* organoids measured in mitochondria stress test by Seahorse analyzer (mean± SEM, n=3, Holm-Sidak’s multiple comparisons test). **c.** Representative images of mitochondria of mt-roGFP2-*shFoxo1/3* organoids. Redox ratio and sum of of oxidized (ex405 em495-590) and reduced (ex488 em495-590) roGFP2. Mean ± SD of 9 crypts control and 8 crypts doxy treated, from 3 independent experiments (For details see materials and methods). **d.** P38 and phosphor-p38 11 levels in doxycycline and Mdivi-1 treated *shFoxo1/3* organoids analysed by western blot, representative of n=3 experiments.

**Suppl. Fig. 3:**
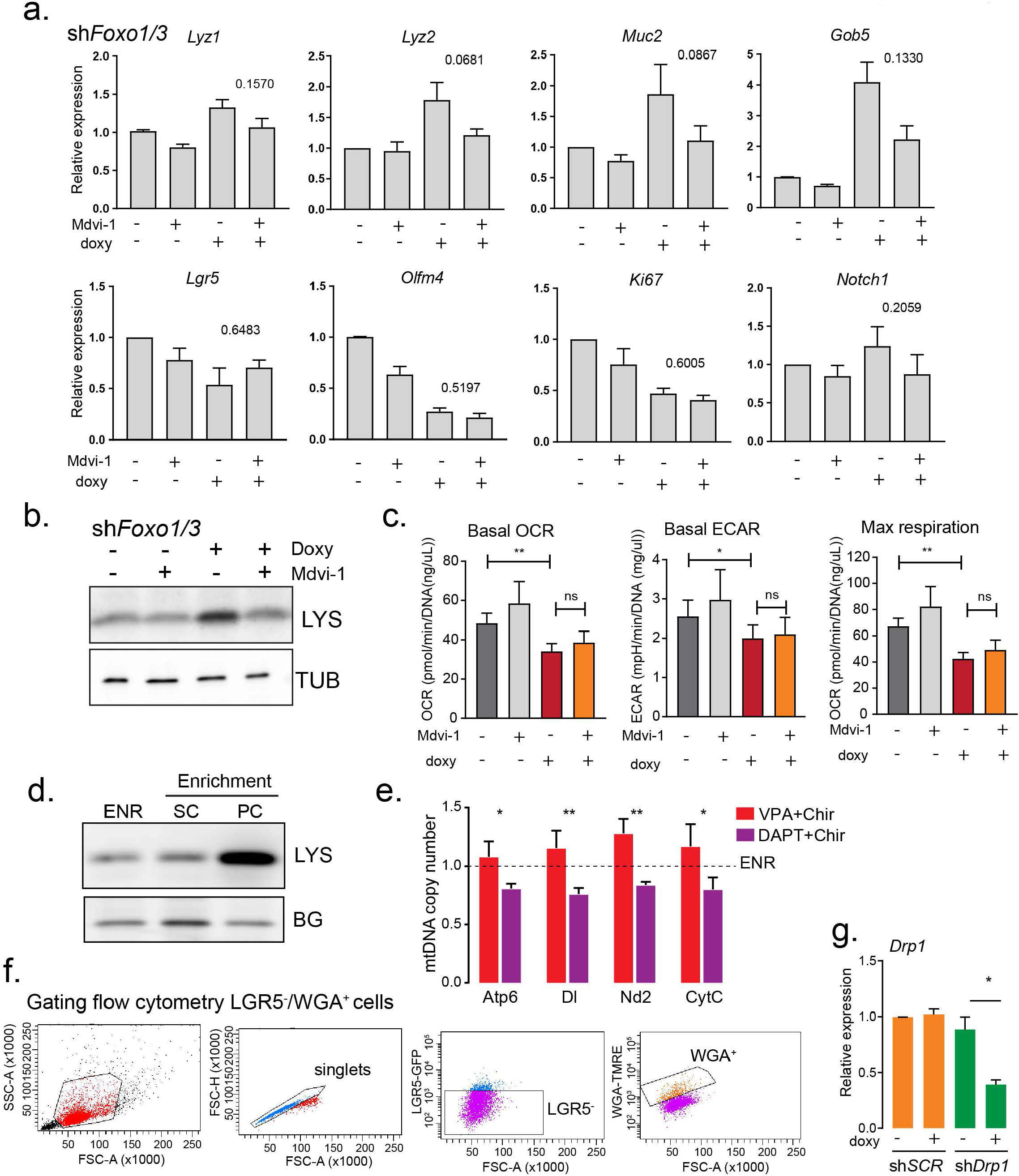
Inhibition of mitochondrial fission partially recovers *Foxo1/3* knockdown-induced secretory differentiation but does not affect bioenergetics. **a.** Relative gene expression in *shFoxo1/3* organoids treated with doxycycline and/or Mdivi-1 determined by qPCR (mean ± SEM, n=5 Kruskal Wallis and Dunn’s multiple comparisons test). **b.** Western blot analysis of *shFoxo1/3* organoids treated with doxycycline and Mdivi-1 (representative for n=4). **c.** Basal OCR, basal ECAR and maximal respiration determined by Seahorse analysis in doxycycline and Mdivi-1 treated *shFoxo1/3* organoids (mean± SEM n=8) Dunn’s multiple comparisons test. **d.** LYS1 analyzed by western blot. ENR control, DAPT: PC enriched, VPA: Stem cell enriched organoids. Organoids were grown for 48 hrs in normal medium (ENR) or with the addition of CHIR9921 and DAPT or CHIR and VPA (representative for n=3). **e.** Mitochondrial copy number of organoids enriched for PCs or stem cells determined by qPCR and represented as ratio to ENR control organoids (mean ± SEM, n=5, ANOVA and Sidak’s test VPA vs. DAPT). **f.** Flow cytometry gating strategy for quantification of live PCs. **g.** *Drp1* 13 mRNA expression by qPCR (mean and SEM n=4 Mann-Whitney Test) *= p < 0.05, **= p<0.01.

**Suppl. Fig.4:**
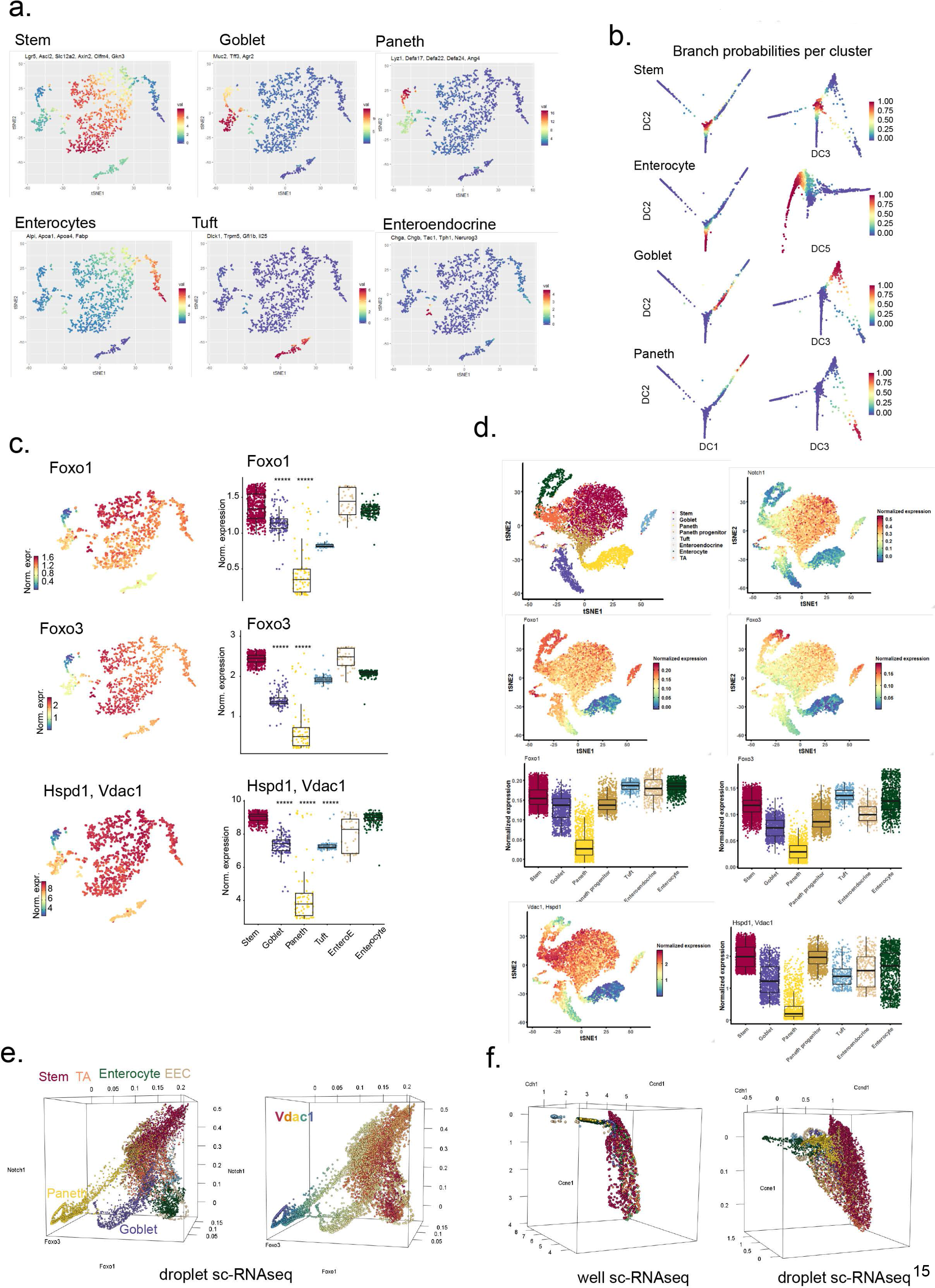
Extended sc-RNAseq analysis reveals a nodal role for Foxo/mitochondria in stem cell differentiation. **a.** Visualization of the mean expression of marker genes for each cell lineage plotted on t-SNE well based/ full-length sc-RNAseq data. **b.** Branch probabilities for each cluster overlaid on the top 1-3 or 1, 2 and 6 DC. Branch probabilities were calculated with Palantir. **c.** *Foxo1, 3* and mitochondrial gene expression levels are plotted on t-SNE plots and boxplots are plotted to represent the quantification of gene expression in each cluster gene. Median values and 25th and 75th percentiles. Mann–Whitney U test, ****= p < 0.0001). **d.** Validation of sc-RNAseq analysis was done by analyzing a second droplet-based sequencing data of 10396 cells (from Harber et al 2017) t-SNE visualization of droplet-based sequencing of Cluster annotation was done on the expression of known cell type markers (same as in a), and Foxo1, 3 and mitochondrial gene expression levels are plotted on t-SNE and boxplots are plotted to represent the quantification of gene expression in each cluster gene. Median values and 25th and 75th percentiles. Mann–Whitney U test, ****= p < 0.0001). **e.** 3 dimensional representation of droplet-based sequencing data gene expression of Foxo1, Foxo3 and Notch1 in intestinal cells. As a fourth dimension cluster classification is assigned by color (left) or colored according Vdac1 expression levels (right). **f.** 3 dimensional representation of well-based scRNA-seq data and second data set of droplet-based scRNA-seq. Genes Ccne1, Cdh1 and Ccnd1 were used as control for comparison with Notch1, Foxo1 and Foxo3. Cluster classification is assigned by color (see c and d).

## STAR★Methods

### Organoid culture

Small intestinal organoids were derived from isolated crypts collected from the entire length of the small intestine of wild-type or DTR–LGR5–GFP mice as described in ref (Sato et al., 2011). The basic culture medium (ENR) contained advanced DMEM/F12 supplemented with penicillin/streptomycin, 10 mM HEPES, 20mM Glutamax, 1× B27 (all from Life Technologies) and 1 mM N-acetylcysteine (Sigma) that was supplemented with murine recombinant epidermal growth factor (Peprotech), R-spondin1-CM (5% v/v, homemade) and noggin-CM (10% v/v, homemade). WENR medium was prepared with ENR medium supplemented with R-spondin1-CM (20% v/v) and 50% WNT3a-CM (conditioned medium) (v/v). A mycoplasma-free status was confirmed routinely. During experiments, organoids were cultured in ENR medium unless stated differently. Treatments were started 36-48 hours after splitting the organoids, depending on when crypt formation was observed. Organoids were treated with 200 ng/ml doxycycline (#D9891, Sigma) for shLuc and shFOXO organoids and 100 ng/ml doxycycline for shSCR, shMFN1/2 and shDRP1 organoids, 25 uM Mdivi (#M0199, Sigma), 5 uM DAPT (#D5942, Sigma), 1 mM Valproic Acid (#P4543, Sigma), 3uM CHIR 99021 (#4423 R&D Systems). Doxycycline and Mdivi-1 treatments on shLuc and shFOXO1/3 organoids were performed for 30-36 hours unless stated differently. CHIR, DAPT and VPA treatments in combination with Mdivi-1 treatments were performed for 48-56 hours.

### Lentiviral transduction

Short hairpin oligos targeting DRP1 (TRCN0000012606) were cloned into Tet-pLKO-puro (#21915, Addgene). Tet-pLKO-puro-Scrambled (#47541) **was** purchased from Addgene. Redox sensor MtGrx1-roGFP2 in pLPCX (Gutscher et al., 2008) was cloned into a lentiviral vector under the control of a Hef1 promotor and with a puromycin resistance cassette. For FOXO1/3 knockdown experiments, doxycycline-inducible pH1tet-flex/FH1tUTG carrying shRNA targeting FOXO1 and FOXO3 (TRCN0000020707) and pH1tet-flex/FH1t(Luc)UTG (Charitou et al., 2015) were used (Charitou et al., 2015). These constructs, together with third generation packaging vectors, HEK293T cells and LentiX Concentrator (Clontech) were used to generate and concentrate lentiviral particles. Organoids were lentivirally transduced as described in (Van Lidth de Jeude et al., 2015).

### Mouse tissue

All procedures involving mice were approved under the protocol 2011-0011 by the institutional animal care and use committee of Weill Cornell Medicine. For the generation of intestine-targeted deletion of Foxo animals Villin-cre allele (JAX# 004586) was introduced to Foxo1/3/4L/L (Paik et al., 2007) by successive breeding.

### DNA, RNA and miRNA extraction and qPCR

For DNA, RNA and miRNA extraction, organoids were washed once with and collected in ice-cold PBS. DNA was extracted with the QIAamp DNA Micro Kit (Qiagen). DNA was used as a template to amplify nuclear and mitochondrial genes. RNA was extracted by using the RNeasy kit (Qiagen), with on-column DNase treatment (Qiagen). cDNA synthesis was performed by using the iScript cDNA synthesis kit (Biorad). MicroRNA extraction and cDNA synthesis were performed by using the MystiCq microRNA kits. QPCR was performed using FastStart SYBR Green Master mix (Roche) and primers listed in table 1.

**Table 1.**
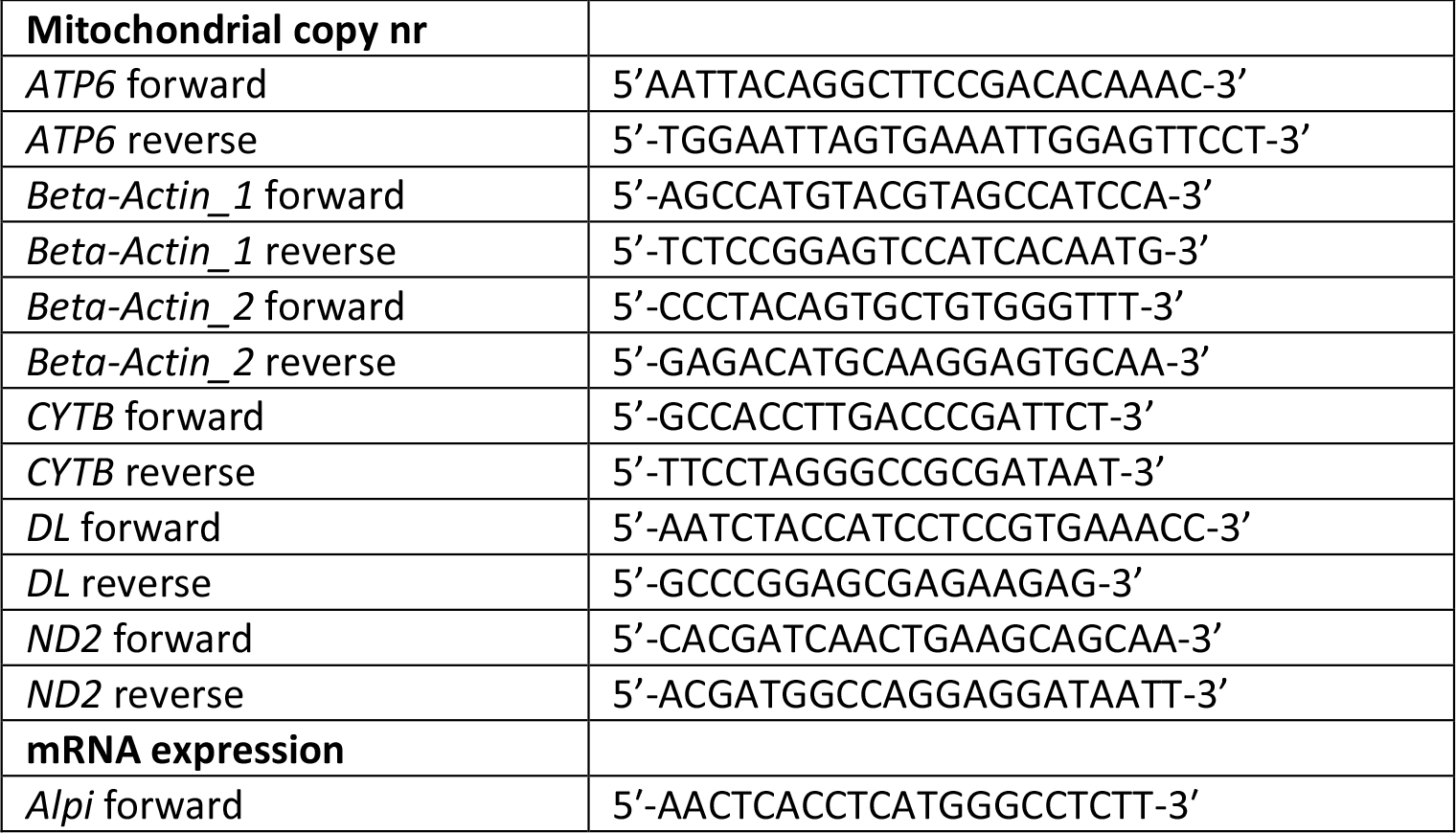
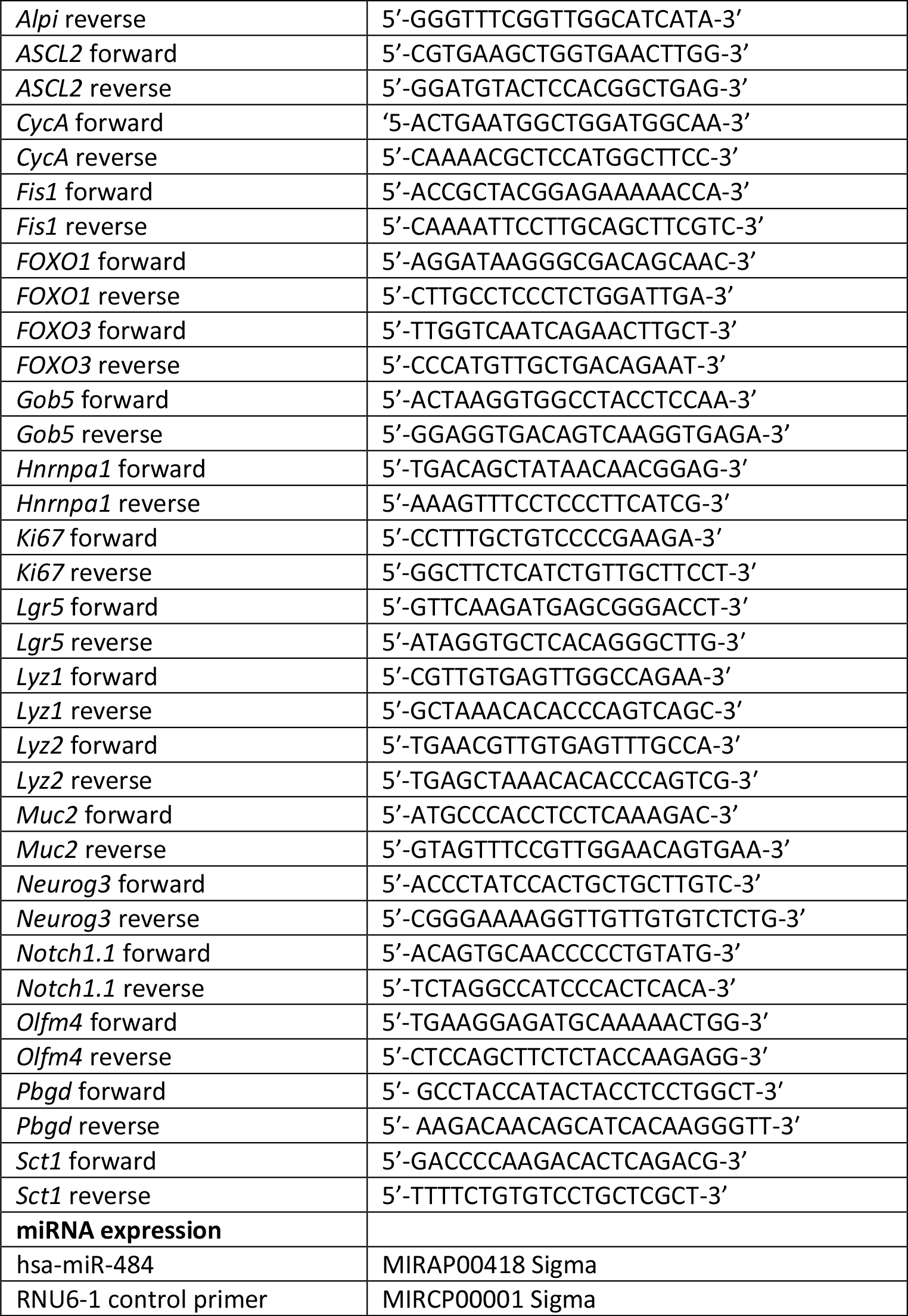
Primer sequences qPCR.

### Protein lysates and western blot

Organoids were washed once with and collected in ice-cold PBS. Total proteins were collected by direct lysis of organoids in Laemli sample buffer. Proteins were run in SDS-PAGE and transferred to Immobilon Polyscreen PVDF transfer membranes (#IPVH00010, Merck Millipore) or Amersham Protan nitrocsllulose membranes (#10600001 GE Healthcare Life Sciences). Western blot analysis was performed with primary antibodies recognizing: GAPDH (Millipore MAB374), vinculin (Sigma V9131), FOXO1 (Cell Signaling 2880), FOXO3 (Novus Biologicals NB100614), lysozyme (Dako A0099), Tubulin (Millipore CP06 OS), Fis1 (SC376469 Santa Cruz), Cleaved Notch1 (Val1744) (Cell Signaling 4147), OLFM4 (Cell Signaling 39141), Secondary HRP-conjugated antibodies targeting mouse and rabbit IgG were purchased from Biorad.

### Seahorse XF flux analysis

Seahorse Bioscience XFe24 Analyzer was used to measure extracellular acidification rates (ECAR) in milli pH (mpH) per min and oxygen consumption rates (OCR) in pmol O_2_ per min. Organoids were seeded in 3 μl matrigel per well in XF24 cell culture microplates (Seahorse Bioscience). 1 hour before the measurements, culture medium was replaced and the organoids were incubated for 60 min at 37 °C. For the mitochondrial stress test, culture medium was replaced by Seahorse XF Base medium (Seahorse Bioscience), supplemented with 20 mM glucose (Sigma-Aldrich), 2 mM L-glutamine (Sigma-Aldrich), 5 mM pyruvate (Sigma-Aldrich) and 0.56 μl ml^−1^ NaOH (1 M). During the test, 5 μM oligomycin, 2 μM FCCP and 1 μM of rotenone and antimycin A (all Sigma-Aldrich) were injected to each well after 18, 45 and 63 min, respectively. For the glycolysis stress test, culture medium was replaced by Seahorse XF Base medium, supplemented with 2 mM L-glutamine and 0.52 μl ml^−1^ NaOH (1 M). During the test 10 mM glucose, 5 μM oligomycin and 100 mM 2-deoxyglucose (Sigma-Aldrich) were injected to each well after 18, 36 and 65 min, respectively. After injections, measurements of 2 min were performed *in triplo*, preceded by 4 min of mixture time. The first measurements after oligomycin injections were preceded by 5 min mixture time, followed by 8 min waiting time for the mitochondrial stress test and 5 min mixture time followed by 10 min waiting time for the glycolysis stress test. OCR and ECAR values per group were normalized to the total amount of DNA present in all wells of the according group.

### Flow cytometry

Organoids were collected in ice-cold DMEM/F12 medium, supplemented with penicillin/streptomycin, 10 mM HEPES and 1x Glutamax (DMEM/F12 +++ medium) and subsequently incubated with trypleE (Life Technologies) for CD24 staining or trypsin (Sigma) for wheat germ agglutinin (WGA) staining to generate single cells. For CD24 stainings, single cells were stained with anti-CD24-APC (#17-0242-82 eBioscience (1:200)) on ice, protected from light, for 30-45 minutes. Excess of antibody was washed away with PBS and cells were resuspended in PBS with DAPI (#D9564, Sigma). Flow cytometry was performed using a BD FACSCelesta #660345. Dead cells were excluded from analysis by excluding DAPI^+^cells. Paneth cells were identified as DTR-LGR5-GFP^−^/CD24^+^/SCC^high^. For WGA stainings, single cells were fixed in paraformadelyde (PFA) (#1004965000, Merck Millipore) at RT for 10 minutes and permeabilized overnight on ice with 70% EtOH. Single cells were washed with PBS and stained with WGA (Wheat Germ Agglutinin) - TMRE (tetramethylrodamine) (1:200, #W7024 thermofisher) protected from light for 1 hr at RT. Paneth cells were identified as DTR-LGR5-GFP^−^/TMRE^+^.

### Immunostaining, live imaging in organoids and immunohistochemistry

We performed immunostaining as described previously (Rodríguez-Colman et al., 2017). In short, organoids were fixed with 4% PFA for 30 minutes at 4°C and permeabilized with PBS buffer containing 10% DMSO, 2% Triton-X100 and 10g l^−1^ BSA. Primary antibodies were incubated o/n (Lysozyme Dako). Paneth cells were stained with WGA and DNA with Dapi. Live imaging of mitochondria was performed in mtGrx1-roGFP1 or after MitoTracker Deep Red FM (#M22426, thermofisher) and Hoechst (#H1399, Life Technologies) staining following manufacturer’s instructions. Imaging was performed using a SP8 confocal microscope (Leica Microsystems). Light microscopy was performed using Nikon eclipse TS100 microscope.

### Mouse intestine immunohistochemistry

Mouse tissues were removed and fixed in 10% buffered formalin. Paraffin embedded intestinal rolls were cut into 5-μm sections. Sections were deparaffinized and rehydrated through a xylene/ethanol series followed by heat mediated antigen retrieval with citrate buffer (pH 6.0). Immunochemistry was performed using the VECTASTAIN Elite ABC kit according to the manufacturer’s instructions. The primary antibodies used were rabbit anti-FOXO1 (1:100; #2880, Cell Signaling Technology), rat anti-BrdU (1:500, #6326, Abcam) and rabbit anti-lysozyme (1:500, #A0099, DAKO).

### qPCR

Mitochondrial copy number was calculated by using the C_t_ method by normalization of mitochondrial gene abundance to presence of nuclear gene b-Actin. Relative gene expression was calculated by using the Ct method by normalization of expression of genes of interest by average expression of housekeeping genes HNRNPA1, PBGD and CycA. Relative miRNA-484 expression was calculated by the Ct method by normalizing the number of miRNA-484 to the total number of miRNAs. Statistical analysis was performed by using Graphpad Prism 7. First Gaussian distribution of data was tested to next apply parametric or non-parametric statistics. Details of statistics are detailed in figure legends.

### Mitochondrial length

Morphology of mitochondria was analyzed by confocal live microscopy in organoids stained with Mitotracker Deep Red and Hoechst. Due to heterogeneity in the morphology of mitochondria across cell types and organoids, we applied quantitative analysis. After image acquisition, we analyzed mitochondrial morphology by using MiNA Image J plugin (Valente et al., 2017). In brief, we first applied smoothing and adjusted B&C. Then, we selected the single cells of interest. Stem cells were selected by Lgr5+ signal and when possible by adjacency to Paneth cells. Paneth cells were selected by Lgr5− signal, nuclear morphology and granule detection in DIC images. We selected 2 consecutive Z-stacks, we cropped the ROI to generate a new image and applied maximal projection. Next, we applied the MiNA macro with default settings and selected the *mean branch length* for each cell as a proxy for mitochondria morphology. Analysis of mitochondrial morphology was performed in independent experiments, taking several organoid crypts from each experiment to compare at least 6 individual cells per condition. For more details per experiment see figure legends.

### mtGrx1-roGFP2

Imaging of mtGrx1-roGFP was performed in 3 independent experiments. Data analysis was performed in ImageJ according to ref. (Gutscher et al., 2008). In short: images of oxidized and reduced mtGrx1-roGFP2 were converted into 32-bit images, automatic thresholding was performed and oxidized/reduced mtGrx1-roGFP2 ratio was visualized by using the image calculator tool. Ox/Red ratios were quantified by measuring the mean of the ratios in whole crypts and single cells.

### Computational analysis

#### scRNA-seq data processing

Two scRNA-seq datasets were downloaded from Gene Expression Omnibus (GSE92332) data repository (Haber et al., 2017). The first contained full-length sequencing data of 1522 cells (8 mice) and the second one contained droplet-based sequencing of 10396 cells. Data processing was done accordingly to the methods described in Haber et al., otherwise stated. Selection of variable genes was performed as described in (Haber et al., 2017). This resulted in 2576 and 1331 variable genes for the full-length and droplet-based datasets respectively. To created TPM-like values transcript counts were multiplied by 10000 for the full-length dataset and then log transformed by computing log_2_(TPM + 1). Finally, batch correction was performed using *ComBat* (Johnson et al., 2007) as implemented in the R package *sva*. PCA and t-SNE and Clustering were computed as described in Haber et al. Detected clusters were validated by plotting known gene markers for the different cell subtypes. The droplet-based dataset showed two different sub-clusters within the enterocyte and paneth populations that were merged.

#### Data imputation

Data imputation was performed using the *magic* function from the *Rmagic* R package (van Dijk et al., 2018) by calculating the k-nearest neighbor graph using Euclidean distance with the previously selected principal components. Note that imputed data is only used in gene trend computation and data visualization. Dimensionality reduction, clustering and pseudo-time calculations were all done using the non-imputed data.

#### Diffusion maps and pseudo-time analysis

We performed dimensionality reduction on the full-length dataset using the diffusion map approach. The diffusion components were computed using *DiffusionMap* function from the *destiny* R package (Angerer et al., 2016) with a local *sigma* and the parameter *k* was set to 500. Pseudo-time and branch probabilities were calculated using the *Palantir* algorithm (Setty et al., 2019) implemented in python. We used as input for the algorithm the first 4 diffusion components after seeing a significant difference between the DC_4_ and DC_5_ eigen values. We defined as early cell a stem cell based on high expression of stem cell markers. We also defined 5 end states that correspond to the canonical differentiated cell types (enterocyte, tuft, paneth, goblet and enteroendocrine) based on the expression of their respective known marker genes.

#### Gene trends

Gene trends were computed by fitting a generalized additive model (GAM) using the *gam* and <u>*s*</u> functions from the *mgcv* R package ([ERROR: Reference with missing citation data for current style.]; Wood, 2003). Gene trends were calculated using MAGIC imputed data. We used thin plate regression splines as the smoothing functions and the parameter *gamma* was set to 10. Gene trends were calculated using all cells because branch probabilities are used as weights, this means that non-committed cells can contribute to all the trends while fully committed cells have a higher weight on their respective trend. We observed that secretory cells appeared very late in pseudo-time which gave some unexpected modelling results, such as very negative expression values due to heavy extrapolation. To circumvent the issue, 30 artificial cells were added to the dataset that contained the same gene expression of the first 30 cells in the ordered pseudo-time. These cells were given equal branch probabilities to the differentiated states. This approach gave smoother and more reliable gene trends for the secretory cells on the earliest pseudo-times (denoted by dashed lines in the plots).

Fast stable restricted maximum likelihood and marginal likelihood estimation of semiparametric generalized linear models on JSTOR.

